# Nucleated Cell Death by a Single Hole Generated by C5b-9

**DOI:** 10.1101/2020.12.02.408690

**Authors:** Anil K. Chauhan

**Affiliations:** From the Division of Adult and Pediatric Rheumatology, Saint Louis University School of Medicine, 1402 South Grand Blvd. St. Louis, MO 63104

**Keywords:** Complement, Membrane Attack Complex, Cell death, One hole, CD4 T cells, C5b-9.

## Abstract

Cell death is essential for tissue regeneration and removal of pathogenic cells. Immunological cell death is caused by membrane attack complex (MAC) or bacterial perforins. A complete understanding of nucleated cell death or bacterial cell death by MAC is essential for therapeutic interventions. The mechanism of MAC mediated cell death in nucleated cell is controversial. We know show that MAC causes cell death in nucleated cells by creating a large single hole in the membrane. Our results are contrary to the current paradigm that states “MAC kills nucleated cell by multiple hits”. MAC deposition on the cell membrane also trigger clustering of membrane rafts, which will have relevant consequences for enhanced cell signaling.

**Significance Statement:** Proteins of late complement pathway that form membrane attack complex, cause death of undesirable nucleated cells such as lymphocytes. The structure of these proteins and their similarity to bacterial cytolysins has been studied. How C9 pore forming protein assemble with rest of C5b-8 on nucleated cell membrane remains controversial. We thus examined and now show the structures formed by these proteins on nucleated cell membranes. Formation of MAC occurs during complement activation during infection and autoimmunity. Largely, our current knowledge of the membrane pore formation is gained using ghost cells, lipid micelles, and erythrocytes. Understanding of how MAC kills nucleated cells will enhance our ability to design new therapies for selective killing of pathogens.

## Introduction

Membrane attack complex (MAC), also referred to as C5b-9 is formed by assembly of the proteins of complement pathway. MAC is a pore-forming complex of the innate immune response. MAC formation is required to kills microbes, microbe infested cells, undesired and tumor cells. Cytolytic activity of MAC, earlier considered to be an enzymatic activity was later explained based on two models, ‘doughnut’ (or ‘channel’) and the ‘leaky patch model’ (1). Arguments support both of these models under a given set of experimental conditions. However, it still remains ambiguous and controversial as to how MAC kills nucleated cells (2, 3). The current model of cytolytic activity of MAC is derived from erythrocytes, synthetic bilayer membranes, bacteria, and lipid-enveloped viruses (2, 4). Recently, the structure of pore forming proteins and their mechanism at molecular level have been solved (5, 6). However, it remains to be addressed as to how these models fit into physical structure of MAC observed on the cell membrane. In this report, we observed that the MAC assembles on a single site on nucleated CD4^+^ T-cell membrane that show calcium efflux and results in the cell death.

Perforins produced by cytotoxic T lymphocytes cause cell death. They share structure and function with C9 protein (7). Both perforins and C9 generate trans-membrane tubule. These trans-membrane tubules are observed in MAC channels that cause cell death (8). Complement proteins C6, C7, C8α, C8β, and C9, and pore-forming toxins (PFT) produced by pathogenic bacteria are globular proteins that contain a common region designated as the membrane attack complex/perforin (MACPF) domain (9). Short peptides and large globular proteins that are secreted by bacteria are also member of this MACPF domain family of proteins (10, 11). These proteins are unique in terms of their ability to switch from a water-soluble form to a membrane-inserted form (12). Recent studies have suggested a shared common mechanism of pore formation among the MACPF domain proteins and cholesterol dependent cytolysins (CDCs) that are the virulence factor of pathogenic bacteria (5, 13, 14). Even though, high-resolution structure of C8;α-MACPF has advanced our understanding of the nature of the protein domain responsible for cell lysis, however in the membrane pore form it still precludes the understanding of arrangements of the protein in complex (5). We analyzed *in situ* MAC formation on CD4^+^ T-cells using purified complement proteins. Unlike previous reports, our results show that MAC assembles at a single site on cell membrane.

## Results and Discussion

In this study, we visualized the *in vitro* pore formation by MAC on Jurkat cells, a CD4 T cell line and peripheral human naïve CD4^+^ T cells by following fluorescently labeled C9 protein (Fig. 1A).The current accepted paradigm propose that in nucleated cells, MAC insertion follows a multiple-hit kinetics, which implies that MAC generates multiple-holes leading to cell death (15, 16). Structural and functional studies have shown that the MAC lesion, which occur at multiple on the cell membrane is initiated by a single C5b-7 or C5b-8 monomer insertion, that is followed by the polymerization of several C9 molecules (17–19). Previous studies with osmotic protection experiments have reported a 1 to 3 nm pore size created by C5b-9 (20). Contrary, our results show formation of MAC at a single site on nucleated cell membrane and these structures observed were heterogeneous in shape and they vary in size and the size of pore complex grows in size over time in the presence of labeled C9 (Fig. 1C)

**Figure 1:**
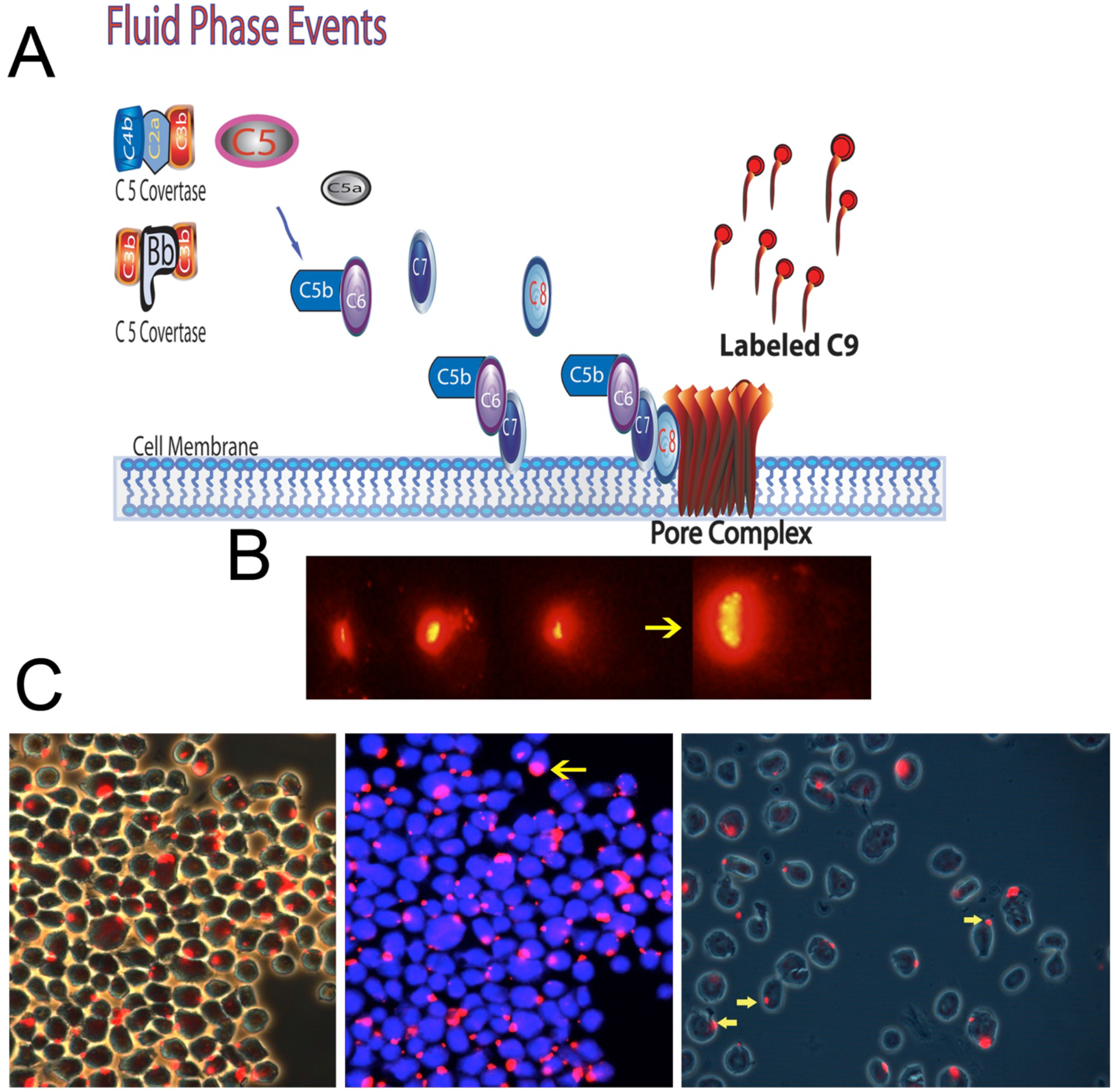
MAC deposits on a single site on CD4+ T cells. (A) Schematic showing the *in-situ* assembly of MAC on Jurkat T-cells. (B) MAC assembly from purified components with labeled C9 show a single site deposit at the cell membrane, at 30 minute (left), 60 minutes (two middle), and 3 h (right). (C) MAC deposits shown on multiple cells, phase contrast (magnification,100X) left, shown same image with DAPI nuclear stain (middle), and at 200X magnification. Yellow arrows point to MAC deposits.

During MAC assembly, surprisingly we observed that all labeled C9 protein polymerized at a single site on the membrane of nucleated cells, occasionally at two sites (Figs. 1 & 4). For control, omission of C8 during MAC assembly or labeled C9 alone did not form these MAC assemblies and no visible staining was observed (not shown). MAC formation occurred as early as within thirty minutes, thereafter these deposits continued to grow in size and formed stable structures on the cell membrane (Fig. 1B). Labeled C9 continued to deposit as observed by the intensity of labeled MAC structures for up to three hours, suggesting an ongoing assembly on the initial single site (right panel on 1B). These results are strange and cannot be reconciled on the basis of a single C5b-8 insertion and subsequent C9 polymerization (C9 of n18) on membrane (21). Based on the size and continuous C9 polymerization in MAC, as evidenced by increase in the size and intensity over extended period of time, we hypothesize that these MAC assemblies could also have been formed from multiple MAC assemblies, which are required to generate a pore size that is effective in causation of cell death (Fig. 1, 2, & 3).

**Figure 2:**
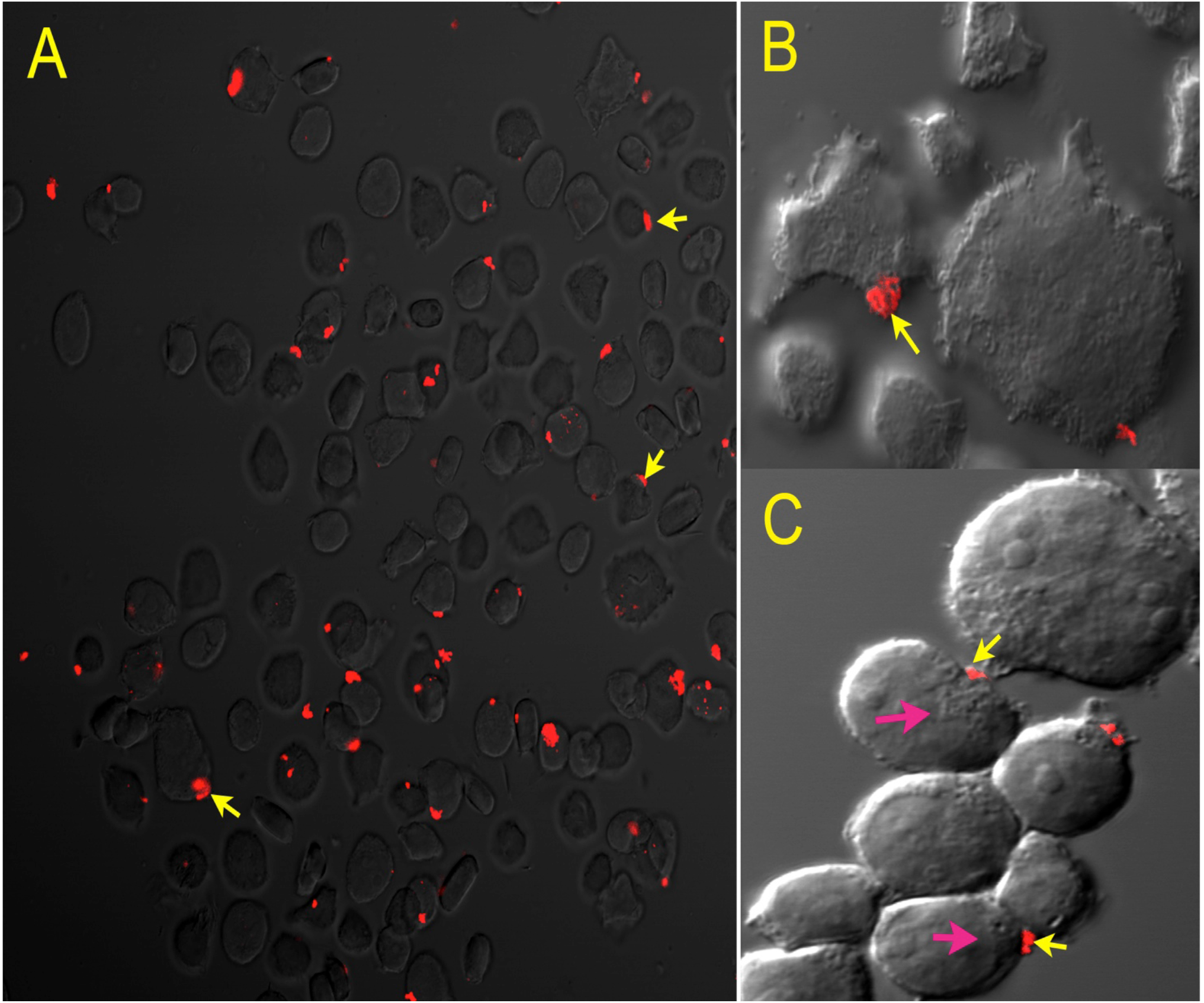
*In-situ* assembled MAC shows heterogeneous structures. (A) Confocal image shows heterogeneity in size and structure of MAC deposits (magnification, 200X) (B) Differential interference contrast (DIC) image showing non-uniform MAC deposit. (C) Confocal image shows dense membrane structures underneath the MAC deposits (magnification, 640X) with 3X optical zoom (pink arrow). Yellow arrows point to MAC deposit.

**Figure 3:**
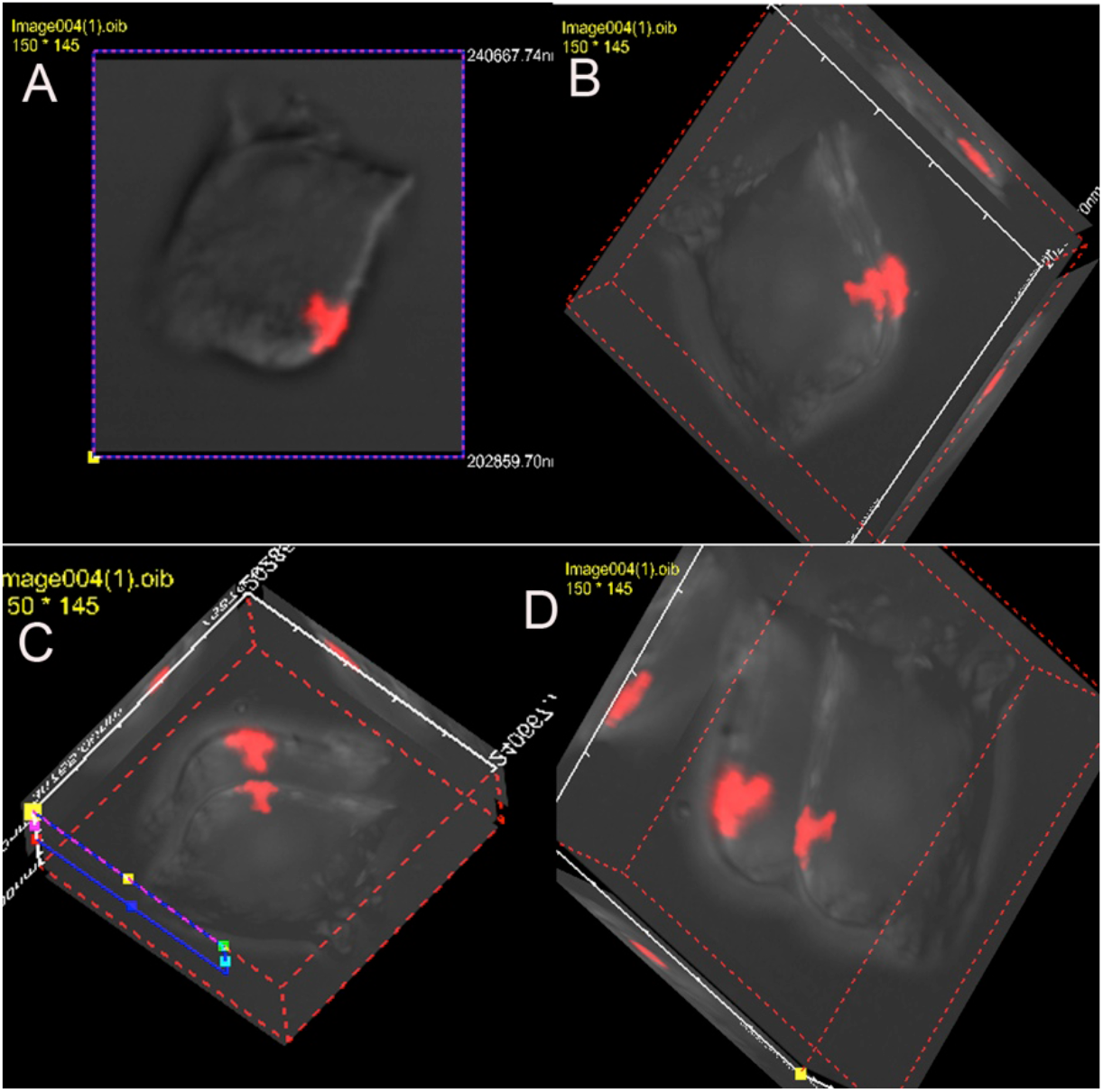
3D view of a MAC deposit. Multiple views of Z-sections upon *in vitro* assembly of MAC from purified C5b-6, C7, C8, and C9-Alexa Fluor 594. A MAC deposit shows a T-shaped structure with most of C9 label on outer membrane, a tubular insertion and flattening of the MAC deposit on inside the cell forming a T-shaped structure.

**Figure 4.**
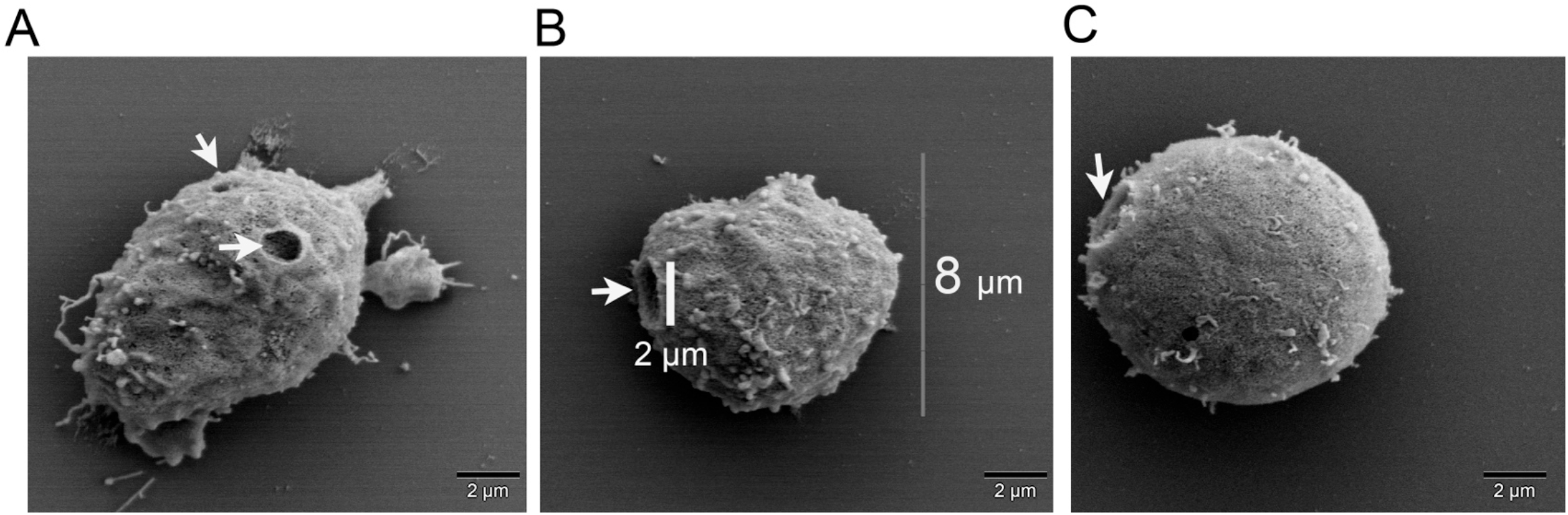
MAC generates a single hole. SEM images of cells post *in situ* assembly of MAC from purified proteins. (A) Naïve CD4^+^ T-cell with two holes. (B) Naïve CD4^+^ T-cell with a single hole. (C) Jurkat cell with a single hole.

These *in vitro* findings do not support the current paradigm of multiple hits hypothesis for nucleated cell death (15). To form the structure observed by us, all multiple hits have to be directed to a single membrane site (Fig. 1 & 2). Alternately, an early C5b-9 deposit on the membrane creates a polarized site, which then further attracts and synchronizes the deposition of additional C5b-7, C5b-8 and/or C9 proteins. In fluid phase, C5b-9 associate with soluble plasma regulators Clusterin or Vitronectin (S protein). Vitronectin bound to C5b-9 is cleaved by granzyme B thereby exposing RGD motif, which then facilitate its attachment to the cell membrane via integrins (22). We assembled MAC in the absence of Clusterin and Vitronectin, which thus exclude a role for Vitronectin in MAC attachment. One has to be careful that these experiments were carried out *in vitro* and may differ from *in vivo* formation of MAC. The presented result does provide an initial observation on as to how a hydrophobic protein can behave on nucleated membrane.

We observed that MAC deposits occupied up to 20% of the T cell membrane. This large size of a MAC deposit cannot be explained by a single protein complex of either (C5b-8)polyC9 or (C5b-8)(2)polyC9 (21). A likely explanation for the observed MAC is that an early single C5b-6 or C5b-7 formation cause the change in the membrane polarity making it conducive for C9 or additional fluid phase C5b-6, C5b-7, or C5b-8 deposition. Alternatively, these MAC structures were formed from coalescence of multiple polymerized C5b-6, C5b-7, C5b-8 or C5b-9 assemblies, originally microscopically invisible on the membrane. These explanations may partially reconcile with the current multiple hit hypotheses, but the cell death likely occur by calcium released from a single pore (Fig. 5) (15). The current model of pore formation by CDC comprises an initial protein attachment to the cell membrane, followed by lateral diffusion and association with other molecules to form a pre-pro oligomer (12). Considering that we were not able to observe these initial assemblies, our results could not be explained by a similar process for the MAC mediated nucleated cell death. The primary site for molecular insertion of the MAC may be a random site or a site that is favored due to the composition of localized lipid content on the nucleated cell membrane. We speculate that membrane rafts (MRs) clustering results in membrane polarization from excessive cholesterol and GM1 gangliosides, contents of MRs. These sites then become the favored site of further deposition of hydrophobic C9 protein or partial MAC complexes.

**Figure 5.**
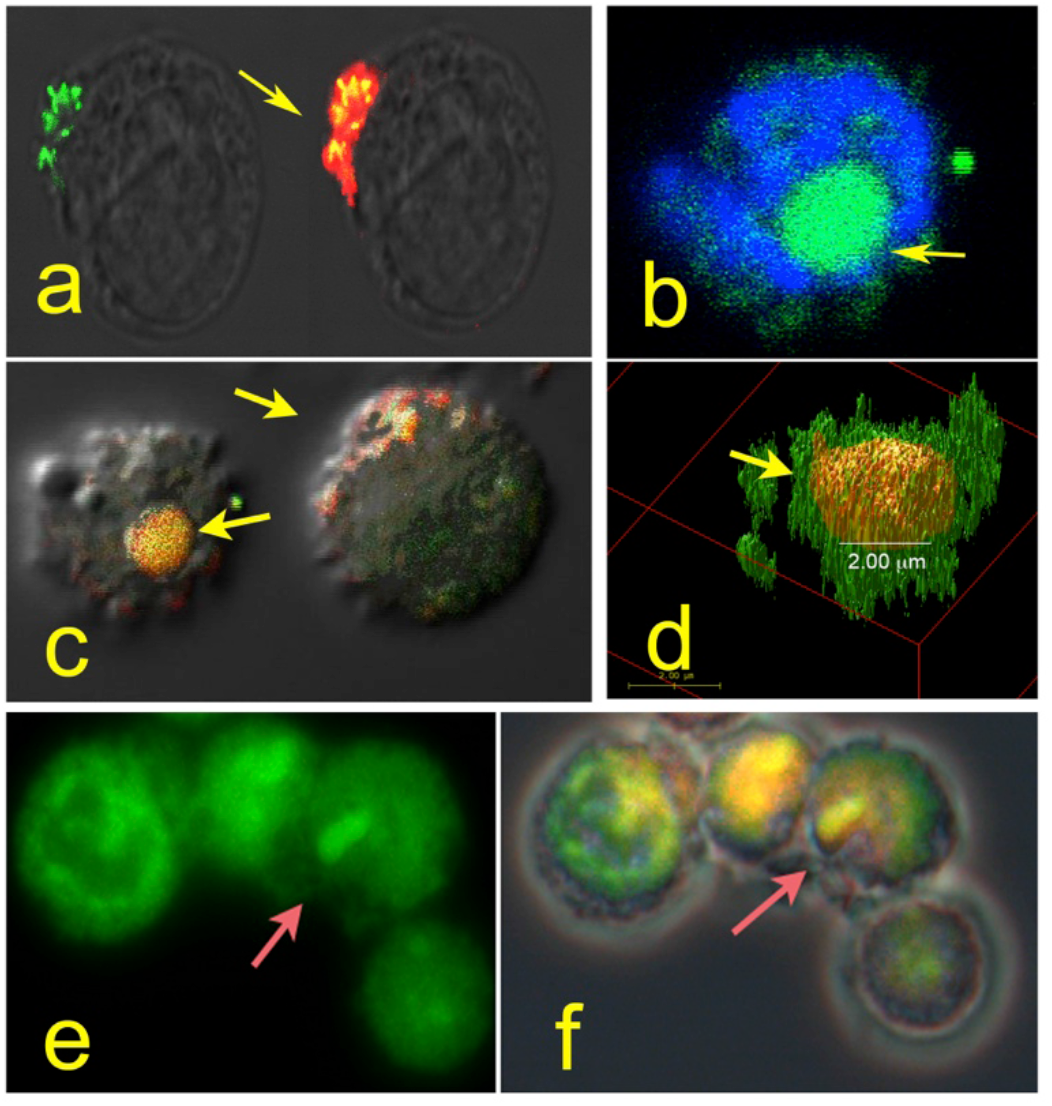
MAC deposit trigger clustering of MRs and show calcium release. MRs (green) forms a cluster underneath MAC deposit (red, a). Calcium is released from a single site (calcium dye, green on DAPI stain, b). DIC image of cell loaded with calcium with *in-situ* assembled MAC show calcium release from single site of MAC deposit (c). A 3D image generated from Z-series sections show ~ 2 μm pore size (d). Microscopy image show cylindrical release of calcium (e) and merge with MAC (f).

Electron microscopic (EM) examination of assembled MAC on the T cell membrane showed a pore size of 2 μm in diameter at a single site and occasionally on two sites in both naïve CD4^+^ T-cells or Jurkat cells, which has an average cell diameter of 11.5 ± 1.5 μm (Fig. 2B & 4) (23). The pore size of 2 μm observed by us is unattainable by a single C5b-9 deposit (Fig. 3). Previous EM studies have shown 18 polymerized C9 proteins in a single pore and proposed that thirty-six; α-helices of C9 are arranged around a circle of 12 nm in diameter (24). These previous EM studies also suggested that C9 undergoes a conformational change from an ellipsoid to an elongated torus (25). Upon polymerization the size of C9 monomer increases from 8 to 16 nm (26). These changes in the protein conformation expose the hydrophobic region of C9, which otherwise is buried within the core of the soluble protein. The amino acid residues 292-333, two predicted amphipathic α-helices of the human C9 map to MACPF region and are suggested to be the key elements required for its insertion into the membrane (27). Our observations do not reconcile with this proposed hypothesis. Another caveat is that this previous work was performed on lipid micelles or ghost cell membranes. The nucleated cell membrane the T cells will have an active state, which will differ in its composition and may not follow a similar kinetic of cell death as observed for erythrocytes or ghost cells.

To further confirm that the cell death occurred by a single hole, we also examined the calcium efflux upon *in vitro* MAC assembly on the cell membrane. Jurkat cells were loaded with calcium binding AA-2 dye, which was followed by in situ MAC assembly. This showed a prominent calcium efflux at 90 minutes from a single site on the cell membrane (Fig. 5 e & f). Both light and confocal microscopy confirmed calcium efflux from a single site. These results further suggest that indeed the nucleated cell death by MAC is caused by the formation of a single membrane pore (Fig. 5).

Our results of a single site MAC deposition do reconcile with the leaky patch model. Previous models of MAC mediated cell death were deduced based on analogical similarities rather than structural information. A mechanistic separation that defines the two earlier models assumes that intrinsic lipids of the target membrane form leaky patches, as proposed in the ‘leaky patch model’, whereas channels are the products of extrinsic complement proteins. *In vitro* MAC deposition showed a single mushroom shaped structure with bulk of the assembly on the outside of cell (Fig. 3A). During an early phase of the complement activation, perhaps the MAC formation starts by forming small random deposits on the membrane, which in sublytic amount are rejected from cell membrane by ectocytosis. In state of hyper complementimia such as in autoimmunity, the rapid assembly of MAC, overcomes the threshold for rejection and promotes coalescence, which finally triggers the pore formation and create an inward protrusion from polymerized MAC on outer membrane (Fig. 3). This structure could reconcile both models by possible generation of single channels on the membranes that eventually coalescence to form a single large patch, as explained by the leaky patch model. Polymerization of hydrophobic MACPF domain results in exposing apolar domains, which results in an increased lipid binding. This hypothesis could possibly also accommodate the biological functions associated with sub-lytic C5b-9, triggering signaling events from bringing required proteins together in MRs (28, 29).

Hydrophobic nature of the assembled MAC structures trigger clustering of MRs underneath them (28). MRs are organelle observed on lymphocyte membranes, which are rich in cytoskeletal and signaling proteins. Staining with fluorescently labeled cholera toxin B (CTB) for MRs in cells with *in situ* assembly of MAC show MRs clustering underneath the MAC deposit (Fig. 5a). The CTB preferentially bind to GM1 gangliosides in the MRs (30). MRs are composed of GM1 gangliosides, cholesterol and other dense lipids and are key participants in lymphocyte cell signaling. MRs appears as dense ring like structures around the MAC deposits (Fig. 2 & 5a). These structures were also observed forming a ring around the pore in scanning EM (SEM) images (Fig. 4). We hypothesize that this apparently occurred from membrane rupture flipping lipids and MRs around the hole. In synthetic bilayer membrane examination using spin-label electron spin resonance (ESR) spectroscopy showed disordered lipids (31). A strong interaction of MAC with lipids of the inner layer of membrane resulted in its partitioning with the MAC during fracturing process affecting 5000 lipid molecules (2). Using electron spin resonance studies, MAC deposits show disordered lipids and lipoproteins (1). Our results provide support to the leaky patch model that contends ‘membrane permeability arise from a local phospholipids’ disorder, reorienting the lipids around inserted apolar protein (31, 32). The lipid-binding capacity of MAC also increases during their assembly. C5b-9 is known to pull lipids out of liposomes, forming extra membranous lipid-protein micelles (32). We thus conclude that these previous studies pointed to lipids are MRs, which upon MAC deposition clustered around and beneath the deposit (Fig. 4A). MR also referred to as lipid rafts, promote oligomerization of; β-PFTs (33).

The soluble fluid phase MAC is shown to trigger functional activity in endothelial cells (28, 34). Since, there is no recognized receptor for MAC, it is likely that these sublytic deposits trigger changes in the membrane potential and more importantly bring the signaling proteins together, a key physiological event for cellular activation in CD4^+^ T-cells (28, 29).

The stability of the MAC on the cell membrane remains controversial. Podack et. al showed destabilization and detachment of MAC bound to protein-lipid micelles (32). Contrary to this observation, two other groups claimed complete stability of liposome-bound C5b-9 complexes (3, 35). We observed stable MAC deposits with MRs on the cell membrane. Occasionally, polymerized MAC detached from the cell membrane. The underlying mechanism that stabilizes MAC association to cell membrane remains ambiguous. We propose that the heterogenic lipid composition influences the overall hydrophobicity of the MAC deposit. Varying C9 content may determine the stability of these structures. The heterogeneity in the size of MAC lesions was observed previously and by us. This has been used as an argument against the channel hypothesis (Fig. 1A, bottom panel). In previous EM studies the ring observed around the MAC was suggested as formed from the excessive C9 content of MAC, while MAC with low C9 appeared as pores without a ring structure (3). We observed a ring like structure in both EM and confocal microscopy. *In situ* MAC was assembled at a molar ratio of 1 (C5b-9): 20 (labeled C9). MAC deposits on CD4^+^ T-cell membrane were heterogeneous in shape and varied in size, even though in SEM images they show a perfect circular pore (Fig. 4). These MAC deposits did not conform to a uniform circular shape reported earlier (Fig. 1 & 2) (36). We observed MAC structures that were also stable during the processing of cells for imaging, which is in agreement with the earlier functional studies where small and large unilammelar vesicles that did not show fusion, fragmentation, or aggregation were observed (35, 37).

In summary, we show that the cell death of human T cell lymphocytes occurs from a single hole created by MAC assembly. In early phase of immune response, low amount of complement activation results in a single site sublytic MAC deposits. These single site sublytic deposits trigger pleiotropic biological responses by forming a synaptic structure that facilitates receptor aggregation that contribute to cellular activation (28). However, the persistent complement activation leads to the formation of a pore, calcium exodus and cell death (Fig. 4 & 5). In small quantities of non-lethal MAC deposits, cells respond by ectocytsis, which further trigger signaling pathways essential for survival such as Akt signaling. However excessive formation of immune complexes (ICs) during infection or tissue damage drives excessive complement activation, which triggers the formation of a pore that cause permeability and loss of calcium. A synchronized role of ICs and C5b-9 in CD4^+^ T-cell differentiation is observed (38, 39). Further studies reconciling the structure of MACPF domain proteins and observation presented in this study should facilitate our understanding of the cell death. Information generated from such studies will be useful in developing agents for selectively killing drug resistant pathogens without causing lymphocyte death. These studies will also provide insight into the mechanisms for selective killing of tumor cells. Cell death in an important process and a better understanding of the underlying mechanisms will advance multiple areas of human health and disease.

## Methods

### Cells and Cell lines

Jurkat cells (J.RT3) were obtained from American Type Culture Collection and maintained as described in the product insert. Human CD4^+^ T-cells were isolated from peripheral blood lymphocytes using naïve CD4^+^ T-cell isolation kit from Miltenyi Biotec (Germany). The blood samples were obtained from Rheumatology Clinic at Saint Louis University with informed consent. These cells were maintained in complete RPMI media with 10% fetal bovine serum, at 37° C and 5%CO_2_, till analyzed.

### Assembly of MAC on cells and Membrane raft staining

A total of 1×10^6^ cells were incubated with purified C5b-6, C7, C8, and labeled C9-Alexa Flour 594. All purified complement proteins were purchased from Complement Tech (USA). The C9 protein was labeled with Alexa Flour 594 using protein-labeling kit (Molecular probe, USA). Cells were washed and starved for 4 h at 37 C. Thereafter, C5b-6 (7 pM), C7 (54 pM), C8 (54 pM) were added and allowed to interact for 10 minutes and then labeled C9 (140 pM) was added. Post formation of MAC assembly, cells were washed with PBS and re-suspended in 0.1%BSA/PBS and a total of 02 μg of cholera toxin B (CTB)-Alexa 488 conjugate was allowed to interact with cells for 20 minute in ice bath. Cells were washed and mounted with anti-fade for examination with regular or confocal microscopy at 400X and 630X magnifications.

### EM studies

Cells after in-situ assembly of MAC using purified proteins were examined in microscopy core using an SEM microscope and images were captured.

### Calcium Release

AA-2 calcium binding dye was loaded in naïve or Jurkat cells as per suggested protocol from vendor. After washing cells were maintained for four hours in plain medium, thereafter, MAC was assembled using purified complement proteins as described earlier. Cells were washed and images were captured using fluorescent microscope and confocal microscope.

## Acknowledgments

Author will like to thank the members of his laboratory for support with the experiments.

